# The wheat cytosolic glutamine synthetase *GS1.1* modulates N assimilation and spike development by characterizing CRISPR-edited mutants

**DOI:** 10.1101/2020.09.03.281014

**Authors:** Yazhou Wang, Wan Teng, Yanpeng Wang, Xiang Ouyang, He Xue, Xueqiang Zhao, Caixia Gao, Yiping Tong

**Author notes:** These authors contributed equally to this work (Yazhou Wang, Wan Teng).

## Abstract

Glutamine synthetase (GS) mediates the first step in the assimilation of inorganic nitrogen (N) into amino acids, however the function of GS encoding genes is not well understood in wheat (*Triticum aestivum*). We found that the cytosolic *TaGS1.1* was the major transcripted *GS1* gene and was up-regulated by low-N availability. CRISPR/Cas9 mediated genome editing was employed to develop two *gs1.1* mutants with mutated *TaGS1.1-6A, −6B*, and *-6D*. Both mutants had lower grains per spike and grain yield per plant than the wild type under both low-N and high-N conditions in field experiments. In a hydroponic culture treated with different N resources, the two mutants was more sensitive to low-N stress than the wild type, but showed similar sensitivity to high ammonium stress with the wild type. The growth deficiency and impaired spike development were associated with the imbalance of N metabolites in the mutant plants. During grain filling, *TaGS1.1* mutation reduced N translocation efficiency and delayed leaf N loss and grain N filling. Our results suggested that *TaGS1.1* is important for N assimilation and remobilization, and required for wheat adaptation to N-limited conditions and spike development.

**Highlight:** The wheat cytosolic glutamine synthetase *TaGS1.1* is important for N assimilation and remobilization, and is required for wheat adaptation to low-N stress and spike development.

## Introduction

Wheat is one of the most important food crops, it alone provides more than 20% of the calories and protein for the world’s population (Braun *et al.*, 2010; Tilman *et al.*, 2011). Breeding and fertilizer application have greatly increased grain yield, and further yield increase is facing the challenges of a slow genetic gain in yield in recent years and efficient use of resources in wheat production (Hawkesford et al., 2013). Early-season nitrogen (N) fertilizer is known to increase tiller/spike number, grain number per spike, whereas late-season N mainly increases the kernel weight and grain protein concentration (Otteson *et al.*, 2008; Peltonen, 1992, 1993). As such, efficient uptake and assimilation of N is critical for the formation of yield components, and it is important to understand the roles of N-use related genes in controlling wheat yields.

Glutamine synthetase (GS) /glutamate synthase (GOGAT) cycle is the first step in the assimilation of inorganic N onto carbon (C) skeletons for the production of glutamine (Gln) and glutamate (Glu). Gln and Glu can then be used to form aspartate (Asp) and asparagine (Asn) through the activity of aspartate aminotransferase (AAT) and asparagine synthetase (ASN) (Coruzzi, 2003). These four amino acids are then converted into all other amino acids and serve as major transport molecules of N between source and sink tissues (Coruzzi, 2003; Galili *et al.*, 2008). There are two GS isoforms in plants, the cytosolic isoform GS1 and chloroplastic isoform GS2. GS1 isoenzymes assimilate ammonium derived from primary N uptake and various internal N recycling pathways (Coruzzi, 2003). GS1 is encoded by a small family of genes that are well conserved across plant species and are crucial for N assimilation and N recycling (Bernard and Habash, 2009). The critical roles of *GS1* genes in N assimilation have been well documented by analyzing N metabolites in the *GS1* mutants of Arabidopsis (*Arabidopsis thaliana*) (Konishi *et al.*, 2017; Konishi *et al.*, 2018; Lothier *et al.*, 2011; Moison *et al.*, 2018), rice (*Oryza sativa*) (Funayama *et al.*, 2013; Kusano *et al.*, 2020; Kusano *et al.*, 2011), and maize (*Zea mays*) (Canas *et al.*, 2010; Martin *et al.*, 2006). The crucial roles of *GS1* genes in N remobilization also have been demonstrated by characterizing the *GS1* deficient mutants (Guan *et al.*, 2015; Kamachi *et al.*, 1991; Masclaux-Daubresse *et al.*, 2010; Moison *et al.*, 2018; Yamaya and Kusano, 2014). *GS1* genes play non-overlapping roles in N use and plant growth. For example, the rice *OsGS1.2* is responsible for the primary assimilation of ammonium in roots, while *OsGS1.1* is important in the process of N remobilization in senescing organs (Funayama *et al.*, 2013; Yamaya and Kusano, 2014). In line with their physiological role in N use, disruption of *OsGS1.2* greatly reduces active tiller number and hence panicle number at harvest; whereas loss-of-function mutation in *OsGS1.1* greatly inhibits rice growth, grain number per panicle, grain size and grain filling (Yamaya and Kusano, 2014). *OsGS1.1* and *OsGS1.2* are unable to compensate for the individual function of another (Tabuchi *et al.*, 2005; Yamaya and Kusano, 2014). The maize GS1 genes *ZmGln1.3* and *1.4* are specifically involved in the control of kernel number and kernel size, respectively (Martin, et al., 2006).

Physiological correlation and QTL mapping have revealed the importance of *GS* genes in N use and yield formation in wheat. GS activity is positively correlated with total N, chlorophyll, soluble protein, ammonium, and amino acids in flag leaves (Kichey *et al.*, 2006; Kichey *et al.*, 2007). N remobilization contributes to approximately 70% of grain N, and this contribution varies among wheat cultivars and closely related to leaf GS activity (Kichey *et al.*, 2006; Zhang *et al.*, 2017c). QTL mapping also has detected the co-localization between QTL for GS activity and N use and yield-related traits (Fontaine *et al.*, 2009; Habash *et al.*, 2007; Li *et al.*, 2015). As such, GS activity can be served as a marker to predict the N status of wheat, and *GS1* genes are considered valuable in breeding with improved N use efficiency (NUE) and yield. For example, overexpression of a *GS1* gene in wheat increased root growth, N uptake, and grain yield (Habash *et al.*, 2001). However, overexpression studies using *GS1* to increase NUE have not yielded consistent results. To develop future strategies for the use of *GS1* in increasing NUE, it is required to understand the pivotal role of *GS1* in the maintenance of essential N flows and internal N sensing during critical stages of plant development (Thomsen *et al.*, 2014). Therefore, understanding the function of *GS1* genes in mediating N use and yield performance will facilitate the use of *GS1* genes in wheat breeding.

Common wheat has three *GS1* genes, one *GS2* gene, one *NADH-GOGAT* gene, and one *Fd-GOGAT* gene in each of the sub-genomes. Our previous studies have shown the essential roles of *GS2* and *NADH-GOGAT* in mediating N use and plant growth in wheat (Hu *et al.*, 2018; Yang *et al.*, 2019; Zhang *et al.*, 2017a). Here we developed *gs1.1* mutants with mutations in *TaGS1.1-6A, −6B*, and *-6D* through genome editing. Investigation of the N use and growth-related traits revealed that *TaGS1.1* is critical for N assimilation, N remobilization, and adaptation to low-N environments. This mutant has reduced grain number per spike and grain yield under both low-N and high-N conditions.

## Materials and Methods

### Plant materials

The winter wheat variety KN199 was used in this study. KN199 was commercially released in 2006, and was used to isolate *TaGS1.1* sequences and develop the *gs1.1* mutants.

### *Genome editing of* TaGS1.1

One sgRNA target for *TaGS1.1* was designed on the conserved domains of all three genomes of wheat variety KN199. The activities of the sgRNA was evaluated by co-transforming the pJIT163-Ubi-Cas9 (Wang *et al.*, 2014) and TaU6-sgRNA (Shan et al., 2013) plasmids into wheat protoplasts. Wheat protoplasts were isolated and transformed as previously described (Shan *et al.*, 2014). The DNAs of plasmids pJIT163-Ubi-Cas9 and pTaU6-sgRNA were simultaneously delivered into immature embryos of KN199 via particle bombardment, as previously described (Zhang *et al.*, 2017b). After bombardment, the embryos were cultured for plantlet regeneration on medium without selective agent. PCR-RE (PCR-restriction enzyme) assays and Sanger sequencing were used to identify wheat mutants in target regions, as described previously (Wang et al., 2014).

### Hydroponic culture

The hydroponic culture was conducted in a greenhouse under the following conditions: 20°C ± 1°C, 50% to 70% relative humidity, 300 μmol photons m^-2^ s^-1^ and a 16-h-day/8-h-night cycle. The germinated seedlings were transferred to the nutrient solution which was described previously (Ren *et al.*, 2012). Three treatments were used, the standard-N (SN), low-N (LN), and ammonium-N (AN) treatments which contained 1.0 mM NH_4_NO_3_, 0.1 mM NH_4_NO_3_, and 4 mM NH_4_^+^, respectively. The nutrient solutions were refreshed every two days.

### Field experiment

The field experiments were carried out in the 2017-2018 and 2018-2019 growing seasons in Hebei Province, China. Both field experiments consisted of two N conditions, each of which had four biological replications. The high-N treatment was applied 18 g m^-2^ N as urea, with 12 g m^-2^ applied before sowing and 6 g m^-2^ applied at the stem elongation stage. The low-N treatment was applied 3.45 g m^-2^ N before sowing. The two N treatments were applied 6 g m^-2^ P as calcium superphosphate before sowing. The seeds were sown in a 2-m-long row with a sowing density of 89 seeds per m^2^. In the 2017-2018 growing season, the seeds were sown in two rows for each genotype in each replicate, and four biological replications were set for each sampling time. At stem elongation, anthesis, 14 days post-anthesis (DPA), and 28 DPA, the aerial parts of five representative plants were collected for dry weight and N analysis. At maturity, the aerial parts of 20 representative plants in each replication were harvested for dry weight, agronomic traits, and total N analysis. In the 2018-2019 growing season, the seeds were sown in four rows for each genotype in each of the four biological replications. At maturity, at least 25 representative plants were harvested in each replication for the measurement of dry weight and agronomic traits. The photosynthetic parameters were measured at 14 DPA by using LI-6400 Portable Photosynthesis System (LI-COR Biosciences, Lincoln, Nebraska USA). Five flag leaves in each replicate were measured in 9-11 am.

### Analysis of N metabolites

The fresh samples stored at −80 °C were homogenized for the measurement of free nitrate and ammonium. The nitrate concentrations in plant tissues were quantified according to the methods described by Cataldo *et al.* (1975), and the free ammonium in plant tissues was determined by Berthelot reaction (Husted *et al.*, 2000). To measure free amino acids, the fresh samples were frozen dried and ground. Ultrasound-assisted extraction was performed for 30 min by adding 1 ml ultra-pure water to 20 mg of ground powder, and the mixture was centrifuged at 10, 000 g for 10 min at 4 °C. The amino acids in the supernatant were derivatized before injection, and then the reaction products were separated and detected by HPLC with a BEH C_18_ sorbent (Waters Alliance e2695, Waters Corporation, Milford, MA). The dried samples were ground for total N analysis by the automated Kjeldahl method (Kjeltec ™ 8400, Foss Analytical A/S, Demark).

### Quantitative Real-time PCR

Total RNA extraction and real-time quantitative reverse transcription PCR (qRT-PCR) were performed according to the methods of Yang et al. (2019). The primers for qRT-PCR were detailed in Supplemental Table S1. The gene expression levels were normalized to the internal control of *TaActin*.

### Western blot and GS activity assay

The fresh plant samples stored at −80 °C were ground to a fine powder under liquid N and then homogenized in an extraction buffer containing 50 mM Tris-HCl (pH 8.0), 2 mM MgCl_2_, 2 mM DTT, and 0.4 M sucrose. The homogenate was centrifuged at 10, 000 g for 20 min two times at 4 °C. The supernatant fraction was used for western blot and GS activity assay. The total protein concentration was determined by a Bradford assay. Western blot analysis was performed using an antibody raised against GS1 protein in rabbits (Abmart, Shanghai, China). The GS activity was determined by using Glutamine Synthetase Detection Kit A047 (Nanjing Jiancheng Biotechnology).

### Statistical analysis of data

Statistically significant differences using SPSS17.0 for Windows (SPSS) were computed based on Student’s *t*-test.

## Results

### *The* TaGS1 *genes differentially respond to N availability*

The published *GS* sequences in wheat were used to blast the reference sequence of Chinese spring in the Ensembl Plants database (http://plants.ensembl.org/index.html). We identified three *GS1* genes and one *GS2* gene in each sub-genome, and the *GS1* genes were named according to their orthologous relation to the *GS1* genes rice (Supplemental Figure S1). The Gene IDs and former names of *GS* genes were presented in Supplemental Table S2.

Gene expression analysis revealed that *TaGS1.1* had much higher transcript abundance than *TaGS1.2* and *TaGS1.3* in roots and shoots of the wheat variety KN199 grown under 1.0 mM NH_4_NO_3_ (SN), 0.1 mM NH_4_NO_3_ (LN) and 4.0 mM NH_4_^+^ (AN) conditions (Fig. 1). Compared with SN treatment, *TaGS1.1* was up-regulated in roots by LN treatment, and down-regulated in shoots by AN treatment (Fig. 1A); *TaGS1.2* was down-regulated by LN and AN treatments in roots, and by LN treatment in shoots (Fig. 1B); *TaGS1.3* was down-regulated in both shoots and roots by LN treatment (Fig. 1C).

**Figure 1.**
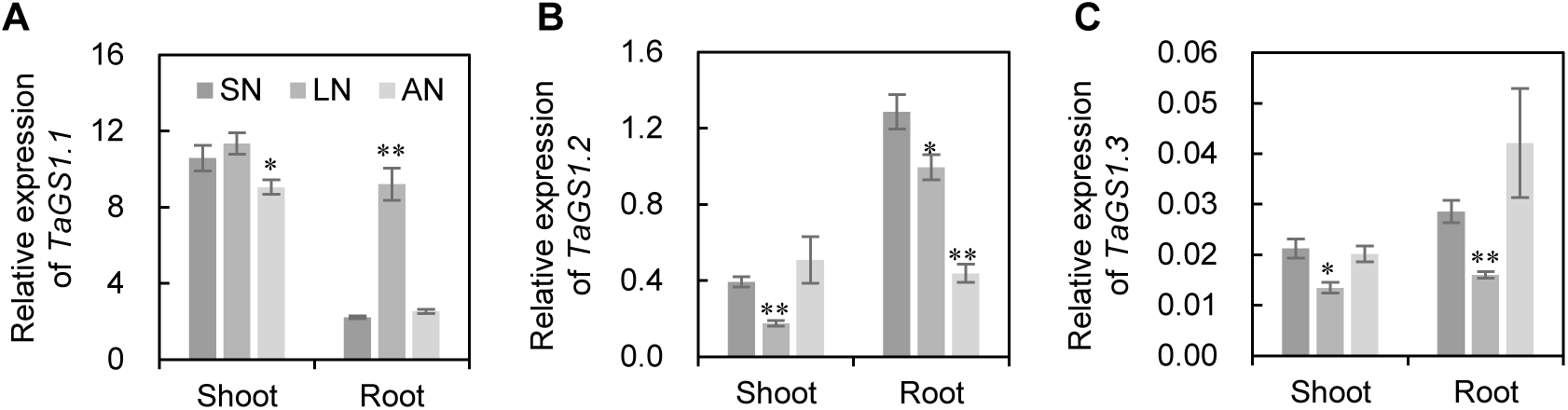
The response of *TaGS1* genes to N availability. A-C, The expression of *TaGS1.1* (A), *TaGS1.2* (B), and *TaGS1.3* (C) in roots and shoots of wheat seedlings grown under 1.0 mM NH_4_NO_3_ (SN), 0.1 mM NH_4_NO_3_ (LN), and 4.0 mM NH_4_^+^ (AN) conditions. Thegerminated seedlings grown for 18 days under SN, LN, and AN conditions. The relative expression levels were normalized to the expression of *TaActin*. The data are expressed as mean ± SE of three replicates. Asterisks indicate the significant difference compared with SN treatment at *P* < 0.05 (*) and *P* < 0.01 (**) level.

We then quantified the expression of *TaGS1* genes in different organs of KN199 grown under field conditions at 20 days post-anthesis (DPA). *TaGS1.1* was most strongly transcribed in leaf blades and sheaths, followed by roots, and the lowest in seeds; *TaGS1.2* was mainly expressed in leaf blades and sheaths, glumes and rachises; whereas *TaGS1.3* was presented at a very low level in all the investigated organs (Supplemental Figure S2A). In flag leaf blades, the expression of *TaGS1.1* displayed a substantial decline from stem elongation (Zadoks growth scale Z37) to 28 DPA, while that of *TaGS1.2* sharply increased from 14 DPA to 28 DPA (Supplemental Figure S2B).

### *Development of* TaGS1.1 *knockout mutants*

Considering the relatively high transcription abundance of *TaGS1.1* and the potential role of *TaGS1.1* in wheat adaptation to low-N availability, we then created *TaGS1.1* knockout mutants of KN199 by using transgenic free CRISPR/Cas9 mediated genome editing. We cloned the three homoeologous *TaGS1.1* genes (*GS1.1-6A, −6B*, and *-6D*) in KN199. A single-guide RNA (sgRNA) that matched perfectly with *TaGS1.1-6A* and *-6D* but had one mismatch with *TaGS1.1-6B* was designed (Fig. 2) in the fifth exon. We used the sgRNA sequence to blast with the Chinese spring reference genomic sequence, and the blast result showed that this sgRNA sequence could avoid off-target (Supplementary Document S1). The genome-editing obtained a mutant. In this mutant, *TaGS1.1-6A* and *-6D* each had homozygous mutation with a 12-bp deletion and a 4-bp deletion respectively, and *TaGS1.1-6B* had heterozygous mutations with a 4-bp deletion in mutation 1 and a 9-bp deletion in mutation 2 (Fig. 2). We grew the T_2_ transgenic plants and WT under high-N and low-N conditions in the 2016-2017 growing season, and identified two kinds of homozygous mutants by PCR-RE (PCR-restriction enzyme) assays and Sanger sequencing. The *gs1.1-1* and *-2* had homozygous mutation 1 and 2 in *TaGS1.1-6B* respectively, and both mutants had homozygous mutations in *TaGS1.1-6A* and *-6B*. Preliminary investigation of agronomic traits showed that both mutants had lower plant height (PH), grain yield per plant (GY) and spike grain weight (SGW) than WT under both high-N and low-N conditions (Supplementary Figure S3A, B, D), but WT and the mutants showed similar spike number per plant (SNPP, Supplementary Figure S3C).

**Figure 2.**
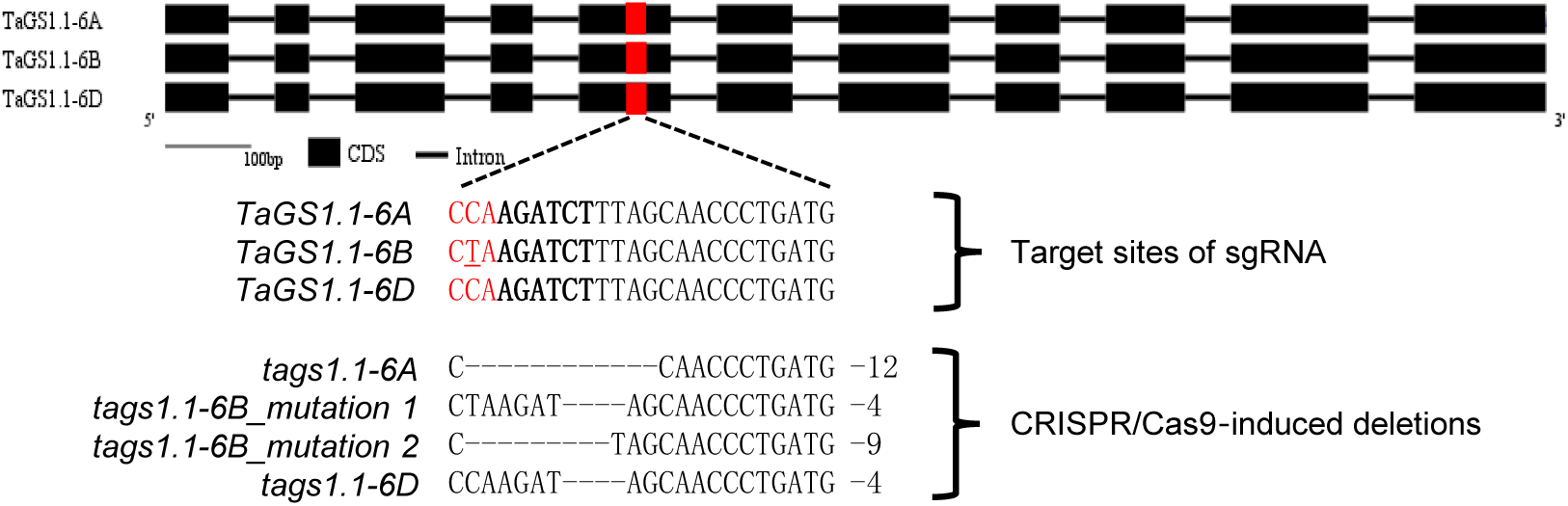
Development of *TaGS1.1* mutant by CRISPR/Cas9-mediated genome editing. Target sites of single guide RNA (sgRNA) in the three *TaGS1.1* homoeologs, and CRISPR/Cas9-induced deletions in the three *TaGS1.1* genes in the *gs1.1* mutants are illustrated. The protospacer-adjacent motif (PAM) is highlighted in red, the single nucleotide polymorphism (SNP) between the three target sites is underlined, the *Bg1*II sites using in the PCR-RE are highlighted in bold font type.

### TaGS1.1 *mutation decreases grain number and grain yield*

We envaluated the agronomic traits in two consecutive growing seasons from 2017-2019 under low-N and high-N conditions. The *gs1.1* mutants headed about two days later than WT (Supplemental Figure S4). The WT and mutant plants did not show a visible difference in leaf and spike color before 14 days post-anthesis (DPA, Fig. 3A, C), but the leaves and spikes of the mutant plants looked greener than those of WT after 25 DPA (Fig. 3B, D). The spikes of the mutant plants looked smaller than those of WT (Fig. 3C, D). Compared with WT in the 2017-2018 growing season, both of the *gs1.1* mutants had a lower GY under high-N and low-N conditions (Fig. 3F). Investigation of yield components showed that the two *gs1.1* mutants had significantly lower spike grain number (SGN) and spike grain weight (SGW) than WT (Fig. 3I, J), but had similar SNPP and 1000-grain weight with WT (Fig. 3G, H). We investigated the morphological traits of the main spikes and found that both of the *gs1.1* mutant has shorter spike length and fewer spikelet number than WT (Fig. 3K, L). We grew the WT and *gs1.1-1* mutant again in the 2018-2019 growing season, the *gs1.1-1* mutant plants exhibited very similar phenotypes in agronomic traits with those in the 2017-2018 growing season (Supplemental Table S3). These two growing seasons clearly showed that the lower GY of the mutant plants mainly resulted from a smaller spike size, which was reflected by spike length, spikelet number, and SGN.

**Figure 3.**
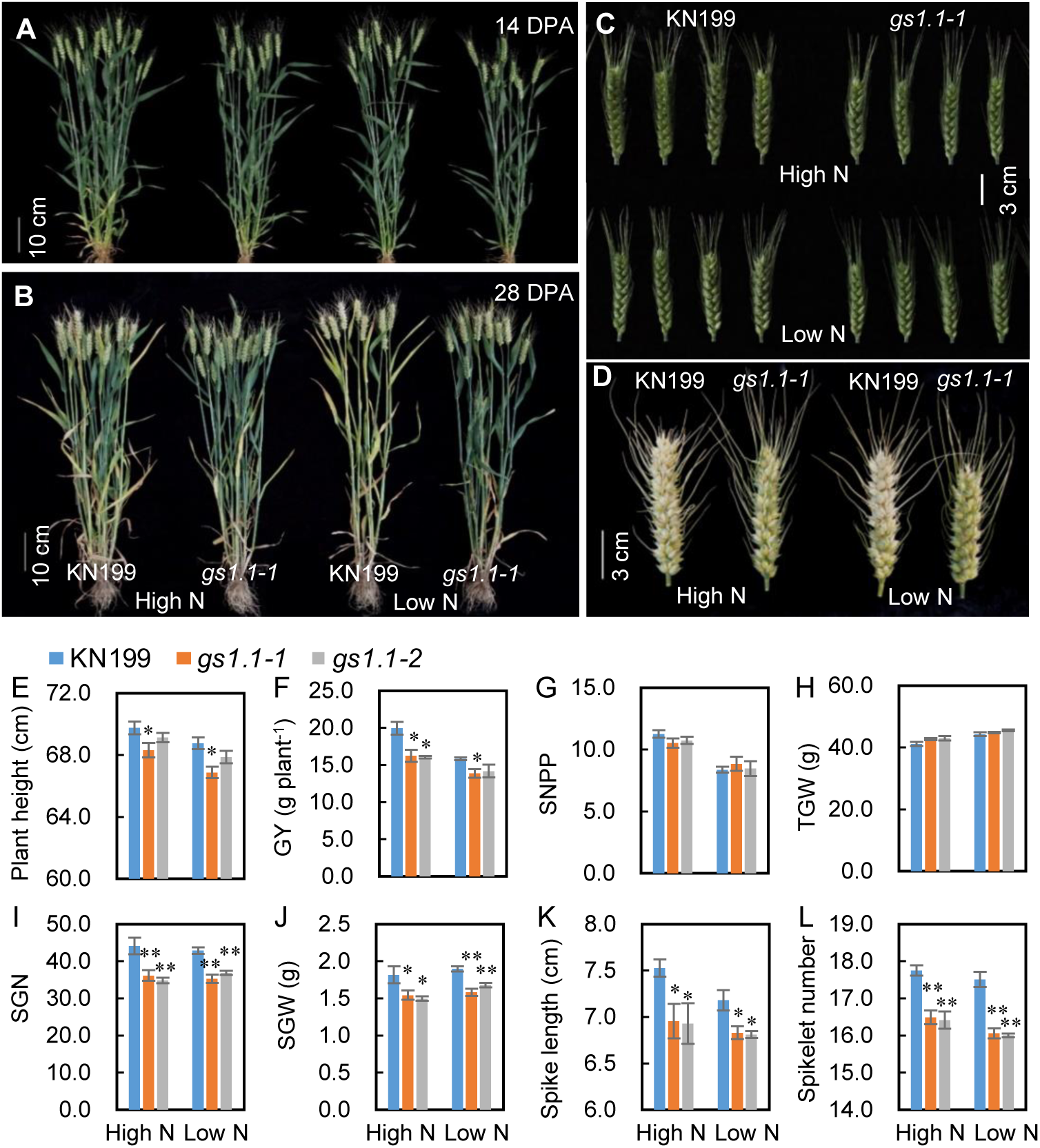
Agronomic traits of the wild type and *gs1.1* mutants under low N and high N conditions in the 2017-2018 growing seasons. A and B, images of plants at 14 days post-anthesis (DPA) (A) and 28 DPA (B), bar = 10 cm; C and D, Spikes at 7 DPA (C) and 30 DPA (D), bar = 3 cm; E, Plant height; F, Grain yield per plant; G, Spike number per plant; H, 1000-grain weight; I, Spike grain number; J, Spike grain weight; K, Spike length; L, Spikelet number per main spike. Data are mean ± SE of four replications. Asterisks indicate statistically significant differences between wild type and *gs1.1* mutants at *P* < 0.05 (*) and *P* < 0.01 (**).

We monitored the dry weight (DW) of different aerial organs in the main culm of the WT and *gs1.1-1* mutant plants at stem elongation, anthesis, 14 DPA, 28DPA, and maturity in the 2017-2018 growing season. All the investigated organs and the whole culm displayed lower DW in the mutant than in WT under both high-N and low-N conditions (Supplemental Figure S5). The relative difference between the WT and *gs1.1* mutant was evaluated by using the DW ratio of the mutant over WT (*gs1.1-1*/WT ratio). The lowest *gs1.1*/WT DW ratio was observed for the spike, the youngest developing organ at stem elongation (Supplemental Figure S5). At stem elongation, the *gs1.1*/WT DW ratio increased with the organ age (Table 1). These results suggested that the inhibitory effect of disrupting *TaGS1.1* on organ DW depended on the organ development stage, the younger the organ, the stronger the inhibition.

**Table 1.** Dry weight for different aerial organs of the main culm at stem elongation in the 2017-2018 growing season.

| Organ | High-N treatment |  |  | Low-N treatment |  |  |
| --- | --- | --- | --- | --- | --- | --- |
|  | KN199 | *gs1.1-1* | *gs1.1-1*/KN199 | KN199 | *gs1.1-1* | *gs1.1-1*/KN199 |
| Spike (mg) | 77 ± 5 | 54 ± 6* | 70.1% | 54 ± 3 | 34 ± 6* | 63.0% |
| Stem (mg) | 413 ± 19 | 350 ± 17* | 84.7% | 396 ± 11 | 334 ± 29 | 84.3% |
| Leaf sheath (mg) | 452 ± 24 | 372 ± 17* | 82.3% | 402 ± 14 | 343 ± 28 | 85.3% |
| Flag leaf blade (mg) | 122 ± 5 | 106 ± 1* | 86.9% | 120 ± 12 | 93 ± 2 | 77.5% |
| Middle leaf blade (mg) | 260 ± 5 | 213 ± 6** | 81.9% | 263 ± 10 | 211 ± 8** | 80.2% |
| Bottom leaf blade (mg) | 133 ± 7 | 123 ± 6 | 92.5% | 136 ± 9 | 128 ± 14 | 94.1% |

Data are mean ± SE of four replications. Asterisks indicate statistically significant differences between wild type and *gs1.1-1* mutant at $P < 0.05$ (\*) and $P < 0.01$ (\*\*).

### *Knockout of* TaGS1.1 *causes root and shoot growth deficiency in an N resource-dependent manner*

Considering the potential role of *TaGS1.1* in mediating N assimilation, we then investigate how *TaGS1.1* mutation affected root and shoot growth of wheat seedlings supplied with different N resources. When the seedling were grown under SN, LN and AN conditions, the *gs1.1-1* and *-2* mutants displayed shorter PH and leaf length, and lower shoot dry weight (SDW), root dry weight (RDW) and root/shoot ratio (DW R/S ration) than WT, in an N resource-dependent manner (Fig. 4). Among the three N conditions, the mutants showed stronger phenotypes in SDW, RDW and DW R/S ratio under LN conditions than under SN and AN conditions, indicating the role of *TaGS1.1* in wheat adaptation to low-N availibility.

**Figure 4.**
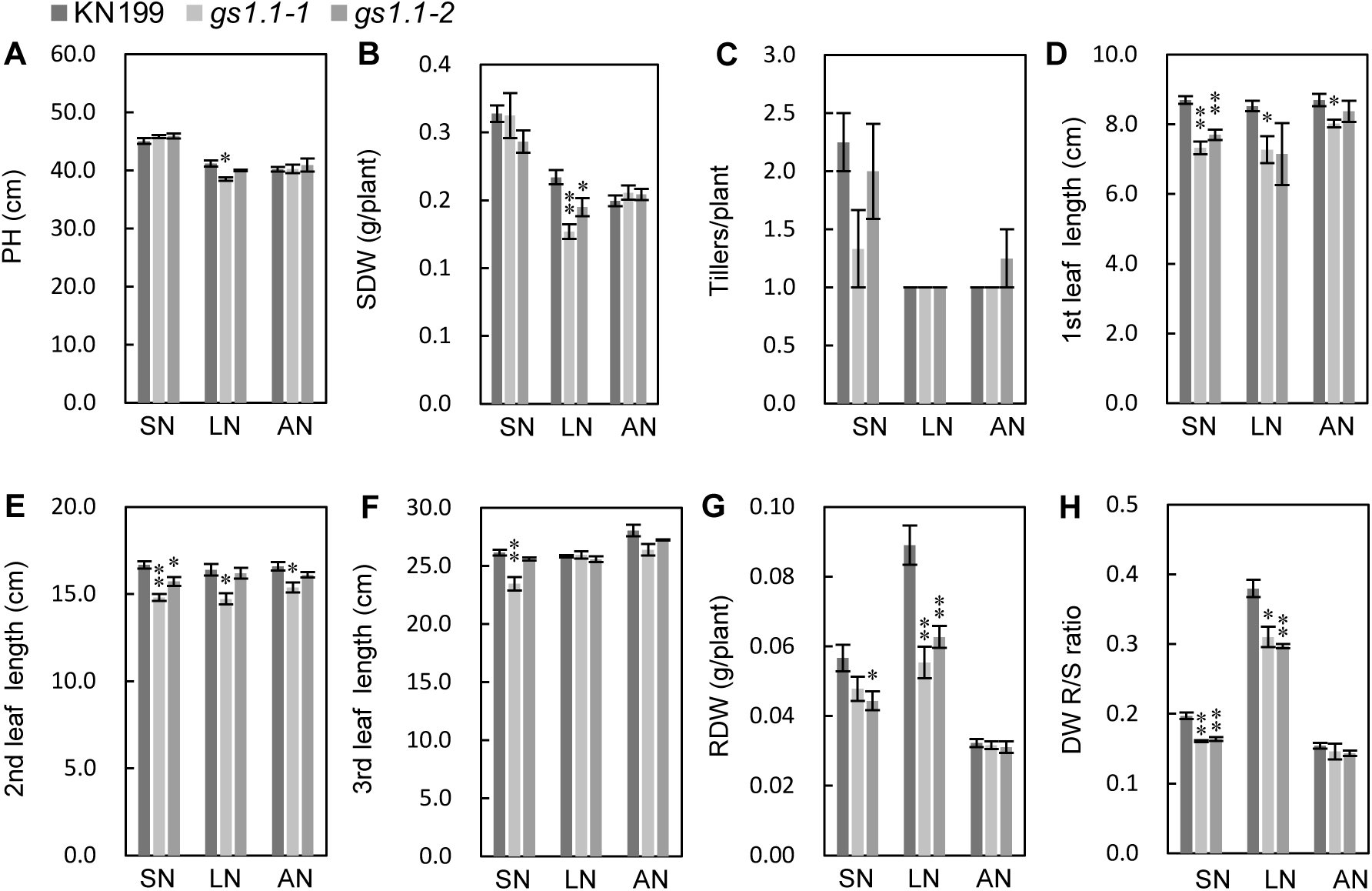
Root and shoot growth-related traits of the wild type and *TaGS1.1* mutants at seedling stage. The germinated seedlings were grown for 18 days under 1.0 mM NH_4_NO_3_ (SN), 0.1 mM NH_4_NO_3_ (LN) and 4.0 mM NH_4_^+^ (AN) conditions. A, Plant height; B, Shoot dry weight per plant (SDW); C, Tiller number per plant; D-F, Length of the 1^st^, 2^nd^ and 3^rd^ leaf; G, Root dry weight per plant (RDW); H, Root/Shoot dry weight ratio. Data are mean ± SE of four replications. Asterisks indicate statistically significant differences between wild type and *gs1.1* mutants at *P* < 0.05 (*) and *P* < 0.01 (**).

### *Knockout of* TaGS1.1 *reduces GS1 protein abundance and GS activity*

To understand the mechanism underlying the growth deficiency of the *gs1.1* mutants, we characterized the GS activity, N assimilation and remobilization of the *gs1.1-1* mutant plants. The seedlings harvested in the hydroponic culture were used to measure these traits. Analysis of gene expression found that the *gs1.1-1* mutant exhibited much lower expression of *TaGS1.1-6B* and *-6D* in roots and shoots than WT under both SN and LN conditions; whereas it showed a comparable expression of *TaGS1.1-6A* with WT (Fig. 5A). It has been reported that a single bp deletion induced frameshift and premature stop prevents mRNA accumulation and consequently results in a low mRNA level of the mutated gene in tobacco (Voelker *et al.*, 1990). The *gs1.1-1* mutant had a much lower GS1 protein level in roots and shoots than WT under SN and LN conditions and lost the LN-induced GS1 protein increase which was observed in WT (Fig. 5B, C). These results suggested that TaGS1.1 was the major GS1 isoform in roots and shoots under both SN and LN conditions at the protein level and majorly contributed to the LN-induced GS1 protein increase. Compared with WT, the total GS activities of the mutant were significantly reduced in shoots under LN conditions, and in roots under SN and LN conditions, whereas those in shoots and roots under AN conditions were not significantly affected (Fig. 5D). These results suggested that the mutant seedlings were deficient in GS activity in the presence of nitrate.

**Figure 5.**
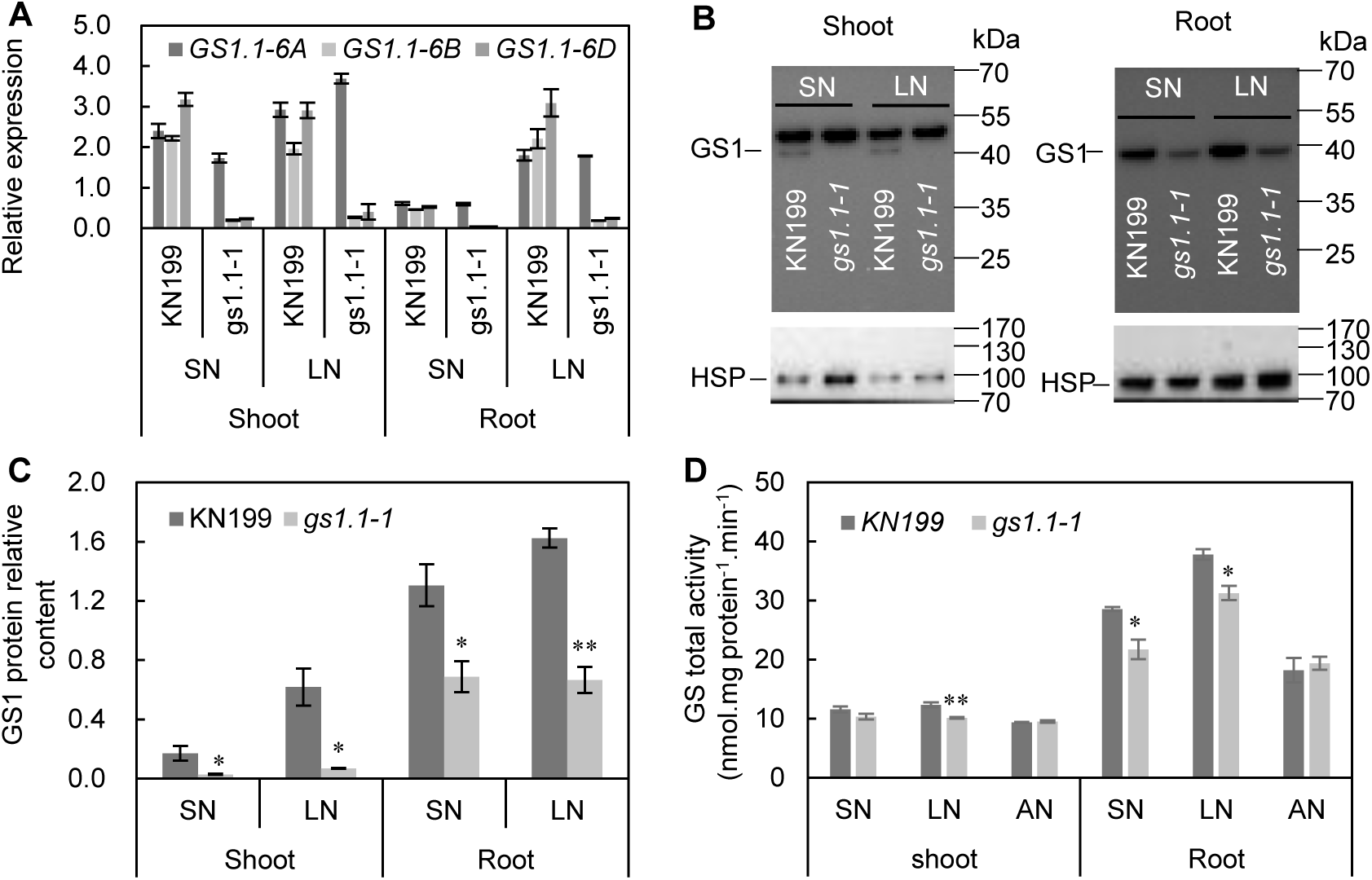
GS activity of the *gs1.1-1* mutant. The germinated seedlings were grown for 18 days under 1.0 mM NH_4_NO_3_ (SN), 0.1 mM NH_4_NO_3_ (LN) and 4.0 mM NH_4_^+^ (AN) conditions. A, Transcriptional expression of the three *TaGS1.1* homeologs in wheat seedling of wild type (WT, KN199) and *gs1.1-1* mutant under SN and LN conditions, the relative expression levels were normalized to the expression of *TaActin*; B and C, Western blot (B) and relative protein abundance (C) of GS1, equal amounts of proteins were loaded in each lane; D, GS activity. Data are represented as means ± SE of four replicates. Asterisks indicate that the difference between the means of the *gs1.1-1* mutant and KN199 was significant at the *P* < 0.05 (*) and *P* < 0.01(**) level.

In the hydroponic culture, the shoots of the *gs1.1-1* mutant plants had significantly lower expression of *TaNR1* (nitrate reductase, Supplemental Figure S6A), *TaNiR2* (nitrite reductase, Supplemental Figure S6B), *TaGS1.2* (Supplemental Figure S6C), *TaNADH-GOGAT* (Supplemental Figure S6E) and *TaFd-GOGAT* (Supplemental Figure S6F) than those of WT, depending on N supply level. In roots, the mutant transcribed significantly fewer transcripts of *TaGS1.2* under SN conditions (Supplemental Figure S6C) and *TaASN1* (Supplemental Figure S6G) than WT under both SN and LN conditions. The *gs1.1-1* mutant and WT had similar mRNA levels of *TaGS2* (Supplemental Figure S6D). We also analyzed the expression of *GS* and *GOGAT* genes in flag leaves in the field experiment and found that knockout of *TaGS1.1* reduced the expression of *TaGS1.1* and *TaFd-GOGAT*, but increased the expression of *TaNADH-GOGAT* (Supplemental Figure S7).

### *The* gs1.1-1 *mutant displays an imbalance of N metabolites*

To understand the roles of *TaGS1.1* in mediating N assimilation, we measured N metabolites in roots and shoots of wheat seedlings exposed to different N recourses as described in Fig. 4. For total N concentration (NC), the WT and *gs1.1-1* mutant only differed in root NC under LN conditions, with the mutant having the lower root NC (Fig. 6A). Compared with WT, the mutant accumulated much higher free NO_3_^-^ in shoots under SN conditions, and roots under SN and LN conditions (Fig. 6B); it also contained significantly higher free NH_4_^+^ in shoots under LN conditions, and in roots under LN and AN conditions (Fig. 6C).

**Figure 6.**
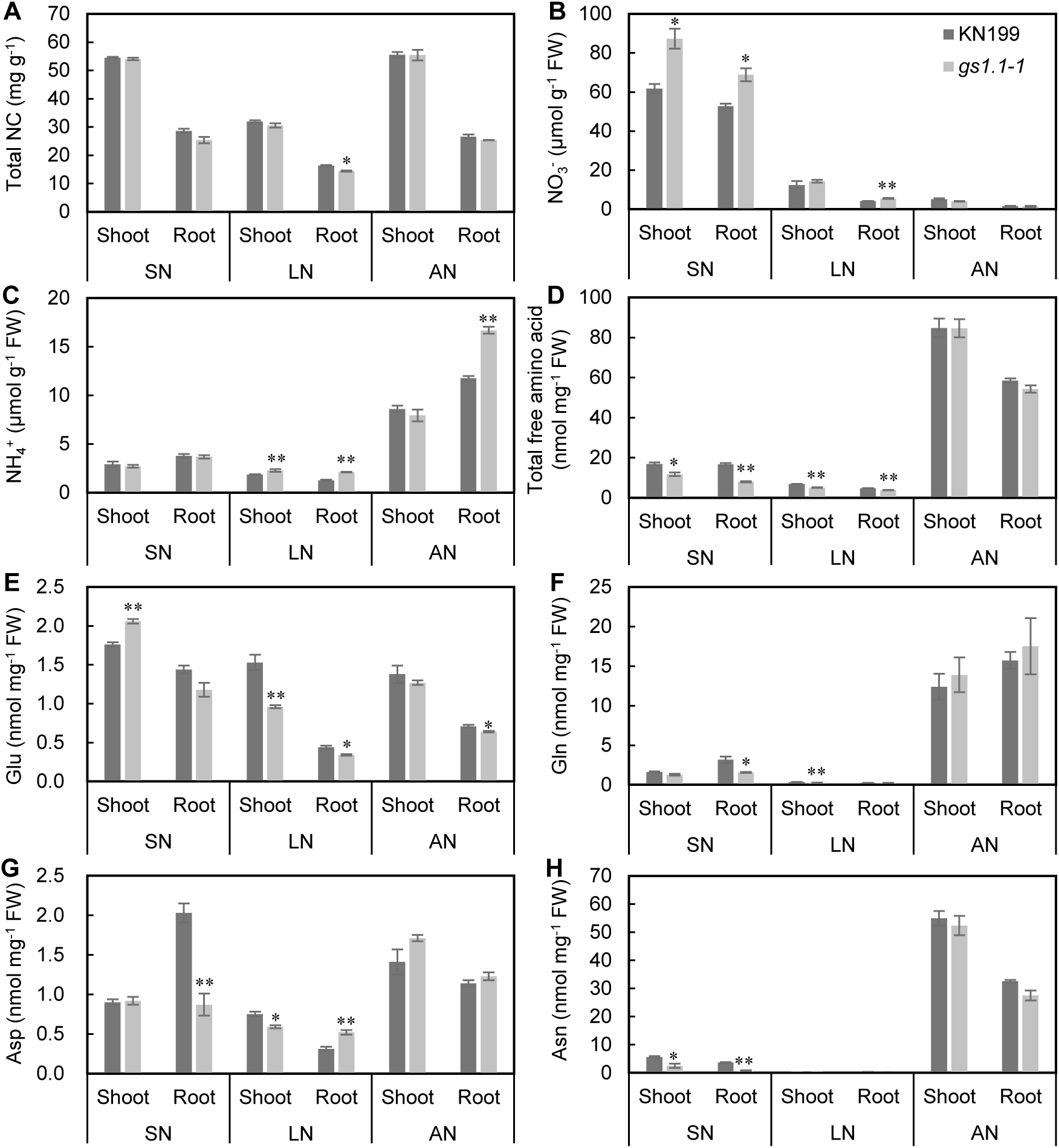
Concentration of N metabolites in roots and shoots of the wild type and *gs1.1-1* mutant. The roots and shoots of wheat seedlings described in Fig. 4 were used to measure the concentrations of total N, nitrate, ammonium, and free amino acids. A, Total N concentration; B, Nitrate concentration; C, Ammonium concentration; D, Concentration of total free amino acids; E, Glu concentration; F, Gln concentration; G, Asp concentration; H, Asn concentration. Data are mean ± SE of four replications. Asterisks indicate statistically significant differences between wild type and *gs1.1-1* mutant at *P* < 0.05 (*) and *P* < 0.01 (**).

The free amino acids were detected in both roots and shoots (Supplemental Table S4). The concentrations of total free amino acids in shoots and roots of the mutant were significantly reduced under SN and LN conditions but not under AN conditions, as compared to WT (Fig. 6D). Under SN conditions, the mutant had significantly lower Asn in shoots, and lower Gln, Asp, and Asn level in roots than WT, but had significantly higher Glu level in shoots than WT; under LN conditions, the mutant had significantly lower Glu, Gln and Asp level in shoots, and lower Glu level in roots than WT, but had significantly higher Asp level in shoots than WT; under AN conditions, the mutant had significantly lower Glu level in roots than WT (Fig. 6E-H). Besides these four amino acids, the levels of many other amino acids were significantly changed in the mutant in a tissue- and N resource-dependent manner, as compared to WT (Supplemental Table S4).

We also measured the free amino acids in the young spikes at stem elongation and seeds at maturity in the field experiment in the 2017-2018 growing season. In the young spikes at stem elongation stage, the concentrations of Asn, Gln, arginine (Arg), alanine (Ala) and proline (Pro) were significantly higher in the *gs1.1-1* mutant than in WT (Supplemental Figure S8A), those of Glu and Asp (Supplemental Figure S8A) and other amino acids (data not shown) did not show significant difference between the mutant and WT. In the seeds at maturity, the *gs1.1-1* mutant had significantly higher levels of Gln and Asn than WT, but had significantly lower level of Asp than WT (Supplemental Figure S8B).

### *Knockout of* TaGS1.1 *decreases flag leaf photosynthetic rate but delays flag leaf senescence*

Since the *gs1.1-1* mutant exhibited growth deficiency, we then investigated if the knockout of *TaGS1.1* affected photosynthetic capacity flag leaves at 14 DPA in the 2017-2018 growing season. The mutant had a significantly lower net photosynthetic rate (*Pn*, Fig. 7A), but significantly higher stomatal conductance (*Gs*, Fig. 7B), intercellular CO_2_ concentration (*Ci*, Fig. 7C); transpiration rate (*Tr*, Fig. 7D) than WT under both high-N and low-N conditions.

**Figure 7.**
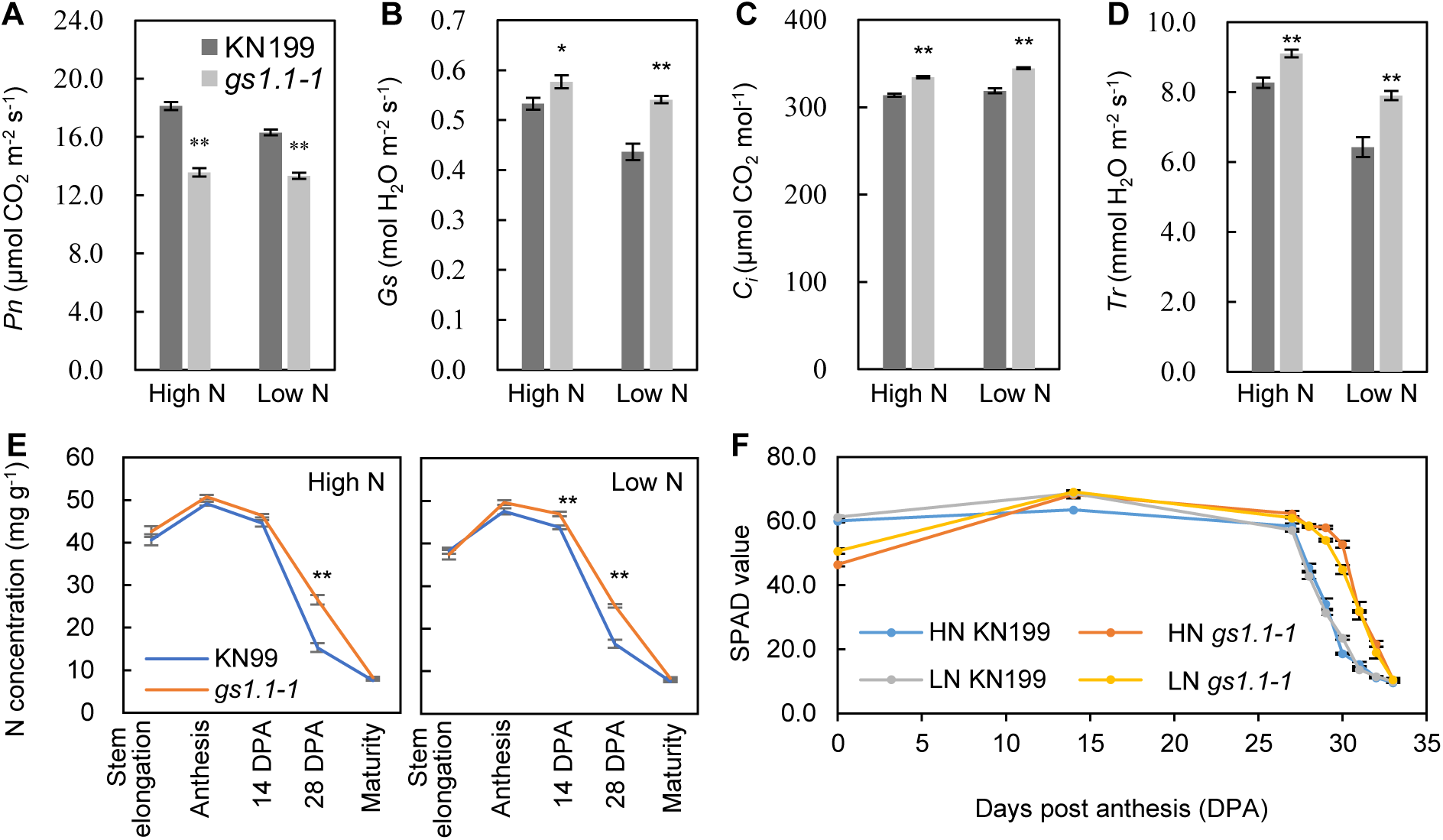
Photosynthetic parameters and N concentrations in flag leaves of the wild type and *gs1.1-1* mutant in the 2017-2018 growing season. A, Net photosynthetic rate (*Pn*); B, Stomatal conductance (*Gs*); *C*, Intercellular CO_2_ concentration (*Ci*); D, Transpiration rate (*Tr*); E, N concentrations in flag leaves; F, SPAD values of the flag leaves. Data are mean ± SE of four replications. Asterisks indicate statistically significant differences between wild type and *gs1.1-1* mutant at *P* < 0.05 (*) and *P* < 0.01 (**).

We also measured total N concentration (NC) in leaf blades, leaf sheaths, stems, spikes (with seeds removed), and grains at stem elongation, anthesis, 14 DPA, 28 DPA, and maturity. The strongest phenotype was observed in flag leaf blades (Supplemental Figure S9C and D), in which the *gs1.1-1* mutant had much higher NC than WT under both high-N and low-N conditions at 28 DPA, but had similar NC with WT at stem elongation, anthesis and maturity stage (Fig. 7E). The dynamic changes in SPAD values clearly showed that the flag leaf senescence was delayed in the mutant as compared to WT (Fig. 7F). These results indicated that *gs1.1* mutation delayed N loss in the flag blades and thereby flag leaf senescence. Similarly, the mutant also displayed a delay of N loss in the stems and middle leaf blades (top 2^nd^ and 3^rd^ leaves) as compared to WT (Supplemental Figure S9A, D). The significantly higher NC in the flag leaf blades and middle leaf blades of the *gs1.1-1* mutant than WT was observed at 14 and 28 DPA under low-N conditions, but was only observed at 28 DPA under high-N conditions (Supplemental Figure S9C, D). The mutant had significantly higher grain NC than WT under both high-N and low-N conditions at maturity (Supplemental Figure S9G).

### *The* gs1.1-1 *mutant has an altered translocation efficiency of N and dry matter during grain filling*

N translocation efficiency (NTE) and dry matter translocation efficiency (DMTE) of aerial organs in the main culm were calculated to reflect N and dry matter remobilization during grain filling by using the data in Supplemental Figure S5 and S9. The *gs1.1-1* mutant had significantly lower NTE in stems, spikes, leaf blades, and sheathes than WT, depending on N supply level (Fig. 8A). The *gs1.1-1* mutant exhibited significantly lower DMTE in stems, leaf sheathes, middle and bottom leaf blades than WT under both high-N and low-N conditions (Fig. 8C). We also calculated N harvest index (NHI) and harvest (HI) and found that the *gs1.1-1* mutant had lower NHI and HI than WT at 14 DPA and 28 DPA, but had similar NHI and HI with WT at maturity under both high-N and low-N conditions (Fig. 8B, D). These results suggested that disruption of *TaGS1.1* delayed grain N and dry matter filling, but not the final NHI and HI.

**Figure 8.**
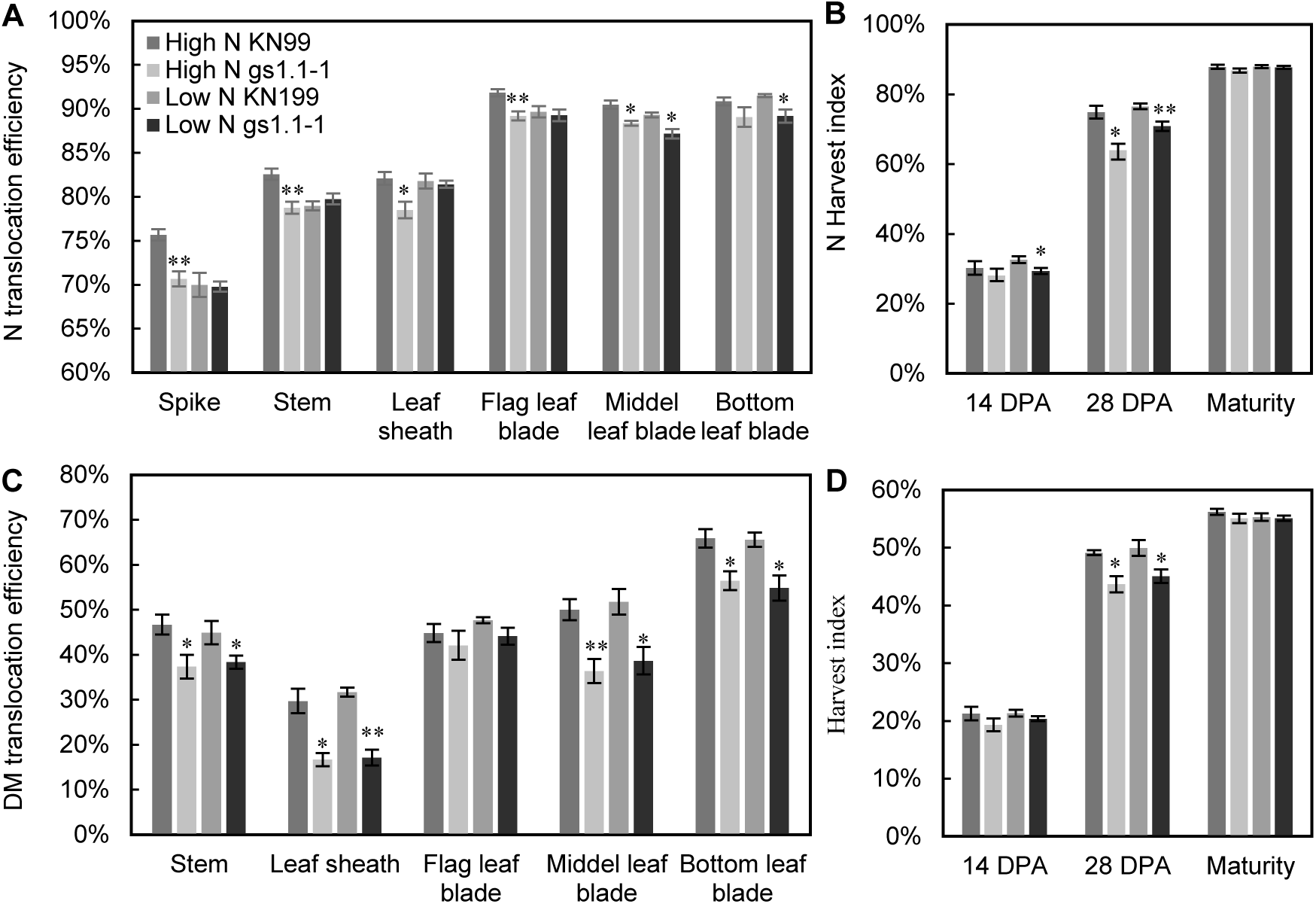
N and Dry matter translocation efficiencies in different aerial parts of the main culm in the wild type and *gs1.1-1* mutant plants in the experiment of the 2017-2018 growing season. A, N translocation efficiency, which was expressed as (N accumulation at anthesis – N accumulation at maturity) / N accumulation at anthesis. B, N harvest index; C, Dry matter translocation efficiency, which was expressed as (dry weight at at anthesis – dry weight at maturity) / dry weight at at anthesis, but the dry matter translocation efficiency for stem was calculated from the data at 14 DPA and maturity. D, Harvest index. DPA, days post-anthesis. Data are mean ± SE of four replications. Asterisks indicate statistically significant differences between wild type and *gs1.1-1* mutant at *P* < 0.05 (*) and *P* < 0.01 (**).

## DISCUSSION

### TaGS1.1 *is the major GS1 isoform*

The expression levels of the three *TaGS1* depended on N availability, organs, and developmental stages (Fig. 1, Supplemental Figure S5), suggesting their non-overlapping roles in N use. The sharp reduction of GS1 protein abundances in roots and shoots of the *gs1.1-1* mutant grown under SN and LN conditions indicated that TaGS1.1 is the major GS1 isoform in both roots and shoots in the presentation of nitrate (Fig. 5C). In line with this result, *TaGS1.1* was more abundantly transcribed than *TaGS1.2* and *TaGS1.3* in shoots and roots at the seedling stage (Fig. 1). A recent study reported that TaGS1.1 has much higher protein abundance than TaGS1.2 and 1.3 in roots and aerial parts (Wei et al., 2019). TaGS1.1 also majorly contributed to the increased GS1 protein level in roots and shoots by LN treatment (Fig. 5C), this result was associated with the up-regulation of *TaGS1.1* by LN treatment, as compared to SN treatment (Fig. 1A).

Although there was a sharp reduction in the GS1 protein level in the *gs1.1-1* mutant, the GS activity was moderately decreased in the mutant under SN and LN conditions, as compared to WT (Fig. 5D). Firstly, a possible explanation for this phenomenon was that the GS activity was measured as the sum of the GS1 and GS2 activities. Secondly, the *gs1.1-1* mutant had four amino acids deletion in TaGS1.1-6A and a frameshift mutation in TaGS1.1-6B and −6D (Fig. 2), the mutated TaGS1.1-6A may still have GS activity. Finally, TaGS1.2 and 1.3 had much higher *V*_max_ for Glu than TaGS1.1 (Wei et al., 2019), the higher *V*_max_ of TaGS1.2 and 1.3 may compensate the effect of low GS1 abundance on GS activity in the *gs1.1-1* mutant. Under AN conditions, the WT and *gs1.1-1* mutant had similar GS activity in both shoots and roots (Fig. 5D). This result was possibly related to the fact that a high level of ammonium greatly reduced TaGS1.1 protein abundances in shoots and roots (Wei et al., 2019). When the external NH_4_^+^ was greater than 2 mM, TaGS1.1 subunit abundance is difficultly detected in shoots and presents at a much lower level than TaGS1.2 in roots (Wei et al., 2019).

### TaGS1.1 *is important for NH*_***4***_^***+***^ ***assimilation***

Measurement of N metabolites revealed the crucial role of *TaGS1.1* in maintenance of internal inorganic N and amino acid homeostasis in wheat plants grown under different N conditions. The results from the seedlings grown in the hydroponic culture, and the young spikes and mature seeds from the field experiment showed that the effects of *TaGS1.1* mutation on N metabolites depended on tissue type and N resource. In the hydroponic culture treated with SN, LN and AN, the *gs1.1-1* mutant had a reduced level of many free amino acids and total free amino acids in shoots and roots under SN and LN conditions compared with WT (Fig. 6, Supplemental Table S4), suggesting the essential role of *TaGS1.1* for NH_4_^+^ assimilation in shoots and roots in the presence of nitrate. However, the *gs1.1-1* mutant had an increased free NH_4_^+^ level in both shoots and roots under LN conditions but not under SN conditions compared with WT (Fig. 6C), indicating that *TaGS1.1* was important for NH_4_^+^ assimilation and homeostasis in wheat plants under LN conditions. The importance of *TaGS1.1* for NH_4_^+^ assimilation under LN conditions was in line with the fact that *TaGS1.1* was up-regulated in roots by LN treatment (Fig. 1A) and majorly contributed to the LN-induced GS1 protein increase in shoots and roots (Fig. 2D). This importance was further supported by a recent study in which TaGS1.1 is found to be expressed in root epidermis cells and displays a higher affinity for Glu and hydroxylamine than TaGS1.2 and TaGS1.3 (Wei et al., 2019). *OsGS1.1*, the rice orthologue of wheat *TaGS1.1* has also been reported its importance in N assimilation under low-ammonium conditions (Ishiyama *et al.*, 2004). Under AN conditions, the *gs1.1-1* mutant had a higher free NH_4_^+^ level and a lower level of Glu and seven other amino acids in roots than WT (Fig. 6C, E; Supplemental Table S4); however, only Ala level was significantly changed in the investigated N metabolites in shoots of the mutant compared with WT (Supplemental Table S4). These results suggested a role of *TaGS1.1* in NH_4_^+^ assimilation in roots, but not (or a limited role, if any) in shoots when NH_4_^+^ was used as the sole N resource. In contrast to the decreased Gln and Asn levels in shoots and roots of the hydroponically grown *gs1.1-1* seedlings in the presentence of nitrate (SN and LN treatments), the field grown *gs1.1-1* plants had higher Gln and Asn in the young spikes and mature seeds than WT (Supplemental Figure S8A). As such, the role of TaGS1.1 in maintaining internal amino acid homeostasis depended on organ type or development stage.

The imbalance of N metabolites in the *gs1.1-1* mutant seemed the combination results of TaGS1.1 deficiency and the down-regulated expression of other genes involved in N assimilation (Supplemental Figure S6). Compared with the shoots of WT, for example, the increased free NO_3_^-^ level was possibly associated with the down-regulated *TaNR1* in the *gs1.1-1* mutant under SN conditions (Supplemental Figure S6A). Knockout of *TaGS1.1* more substantially reduced the total free amino acid level in shoots and roots under SN conditions than under LN conditions compared with WT (Fig. 6D), this was possibly associated with the fact that the transcripts of *TaGS1.2* in shoots and roots of the mutant were significantly reduced under SN condition but not under LN conditions, as compared to WT (Supplemental Figure S6C). Among the detected free amino acids, the Asn level in roots under SN conditions was most sharply reduced (the lowest *gs1.1-1*/KN199 ratio, 0.19) by disrupting *TaGS1.1* (Supplemental Table S4). As ASN catalyzes the synthesis of Asn and Glu from Asp and Gln, the reduced *TaASN1* transcripts together with the reduced Asp and Gln level might contribute to the sharp reduction of Asn in the *gs1.1-1* roots under SN conditions (Fig. 6F, G; Supplemental Figure S6G).

The internal pools of amino acids within plants have been suggested to indicate N status of a plant, and are served as a signal to regulate gene expression (Miller *et al.*, 2008). Compared with SN treatment, LN treatment had a lower level in total NC and most of the investigated N metabolites in both shoots and roots of WT (Fig. 6, Supplemental Table S4), indicating an N-deficient status of WT under LN conditions. The *gs1.1-1* mutant had a reduced level of many free amino acids and total free amino acids in shoots and roots under both SN and LN conditions (Fig. 6D, Supplemental Table S4). As such, the mutant might have a deficient N status under SN conditions and a more deficient N status under LN conditions compared with WT, and thereby displayed growth deficiency. In line with this claim, the responses of several N-assimilation genes to N availability were changed in the mutant. In shoots, the expression of *TaNR1, TaNiR2*, and *TaGS1.2* in WT was up-regulated by SN treatment as compared to LN treatment; however, these genes had significantly lower expression in the mutant than in WT under SN conditions (Supplemental Figure S6A-C). In roots, the expression of *TaGS1.2* and *TaASN1* in WT was up-regulated by SN treatment as compared to LN treatment; however, disrupting *TaGS1.2* impaired the SN up-regulated *TaGS1.1* expression, and significantly reduced *TaASN1* expression under SN and LN conditions (Supplemental Figure S6C, G).

### TaGS1.1 *functions in N remobilization*

Cytosolic GS1 isoforms are known to positively mediate N remobilization during leaf senescence (Moison *et al.*, 2018; Yamaya and Kusano, 2014). Our current study showed that knockout of *TaGS1.1* in the *gs1.1-1* mutant reduced N and dry matter translocation efficiency during grain filling (Fig. 8A, C), and delayed N loss and senescence of flag leaves (Fig. 7E, F). However, disruption of *TaGS1.1* did not significantly affect the final NHI and HI (Fig. 8B, D) and total N concentrations in the vegetative organs at maturity (Supplemental Figure S9). These phenomena were possibly associated with the dynamic changes of *TaGS1* transcripts. In flag leaves, the mRNA levels of *TaGS1.1* and *1.2* sharply decreased and increased after 14 DPA, respectively, and *TaGS1.1* had a higher mRNA level than *TaGS1.2* before 14 DPA, but a much lower level than *TaGS1.2* after 14 DPA (Supplemental Figure S2B). These results indicated that *TaGS1.1* may mainly function during relatively earlier grain filling, while *TaGS1.2* may mainly function during relatively later grain filling in modulating N and dry matter remobilization. Since knockout of *TaGS1.1* did not significantly alter the expression of *TaGS1.2* in flag leaves (Supplemental Figure S7B), it can be assumed that the functional *TaGS1.2* may partially compensate for the adverse effects of *TaGS1.1* deficiency on N and dry matter remobilization.

### TaGS1.1 *is required for wheat tolerance to low-N stress*

The three *GS1* genes showed a differential response to N availability (Fig. 1), suggesting their different roles in wheat adaptation to a fluctuating nutrient environment. In the hydroponic culture, the *gs1.1-1* and *-2* mutant seedlings were more sensitive to low-N stress than WT (Fig. 4). At stem elongation, the young spike growth of the *gs1.1-1* mutant was more sensitive to low-N treatment than that of WT under field conditions (Table 1). The role of *TaGS1.1* in wheat adaptation to low-N stress was supported by its importance for NH_4_^+^ assimilation under N-limited conditions, which has been discussed in the context. The amino acids Glu, Gln, Asp, and Asn are the precursors of other amino acids in N assimilation, and Glu and Gln may serve as signals of organic N status (Gutierez *et al.*, 2008). The reduced low-N tolerance of shoot growth in the *gs1.1-1* mutant was possibly associated with the fact that the mutant had a reduced level of Glu, Gln, and Asp in shoots under LN conditions but not under SN conditions, as compared to WT (Fig. 6E-G).

A high level of ammonium in the growth media is known to inhibit plant growth, as has been shown in our current study (Fig. 4). The WT and *gs1.1* plants did not differ in their sensitivity to high ammonium stress under our experimental conditions (Fig. 4). The underlying reason might be because that *TaGS1.1* had a limited role in NH_4_^+^ assimilation in shoots under AN conditions. The wheat GS1.1 is orthologous to the rice GS1.1/Gln1.1. However, the rice *gs1.1* mutant exhibits a severe decrease in shoot growth and an imbalance in the levels of sugars, amino acids, and metabolites when ammonium is used as the sole N resource (Kusano *et al.*, 2011). Since nitrate and ammonium are, respectively, the major N resources for wheat and rice, it can be assumed that *TaGS1.1* and *OsGS1.1* are evolutionally adapted to nitrate and ammonium recourses, respectively.

### *TaGS1.1* Is Required for Spike Development

We observed plant growth deficiency in the two *gs1.1* mutants at the seedling stage in the hydroponic culture (Fig. 4) and in the field experiments (Fig. 3, Supplemental Figure S3). In line with these results, the *gs1.1-1* mutant exhibited much lower *Pn* in flag leaves than WT under both high-N and low-N conditions at 14 DPA (Fig. 7A). The impaired plant growth and photosynthetic capacity were possibly caused by a deficiency in free amino acids and the imbalance of N metabolites but not by total N deficiency in the mutant. In fact, the mutant had similar with, or even higher N concentrations in aerial parts than did WT (Supplemental Figure S9).

The field experiments showed that the lack of *TaGS1.1* reduced GY by inhibiting spike development, as both of the *gs1.1-1* and *-2* mutants had a shorter spike length, a fewer spikelets and grains per spike, and consequently a lower spike grain weight compared with WT (Fig. 3, Supplemental Figure S3, Supplemental Table S3). Monitoring aerial organ growth of the *gs1.1-1* and WT plants from stem elongation to mature revealed that the growth deficiency in the *gs1.1-1* mutant was most obvious in the young spikes at stem elongation stage (Table 1). This result together with the later heading date in the mutant (Supplemental Figure S4) indicated a delay in spike development in the mutant compared with WT. Calculation of the data in Fig. 3I and 3L showed that the *gs1.1-1* and *-2* mutant plants had fewer spikelets per spike and lower grains per spikelet (the ratio of grain number per spike over spikelet number per spike) than WT, indicating that both of spikelet development and floret development (or seed setting rate) were impaired in the mutants. Measurement of free amino acids in young spikes at stem elongation stage (corresponding to the floret development stage) revealed the over-accumulations of Gln, Asn, Arg, Ala and Pro in the *gs1.1-1* mutant plants (Supplemental Figure S8A), and these over-accumulations maybe associated with the impaired spike growth in the mutant. Maintenance of internal amino acid homeostasis has been shown the importance plant growth and development (Lu et al., 2018). Loss-of-function of *OsARG* encoding an arginine hydrolysis enzyme increases Arg accumulation in panicles and leads to small panicle and low seed-setting rate in rice (Ma *et al.*, 2013). Exogenous application of a low level of Lys, Arg, Val, and Ala to nutrient solution promotes tiller bud outgrowth, but applying a high level of these amino acids inhibits tiller bud outgrowth in rice (Lu *et al.*, 2018). Overexpression of an amino acid permease, *OsAAP5*, in rice transports more Lys, Arg, Val and Ala to the tiller base parts and reduces tiller bud outgrowth, while the opposite result occurs in the RNAi-mediated knockdown lines (Wang *et al.*, 2019). These results suggest that an appropriate concentration range of free amino acids is required for tiller and spike growth. As such, it can be assumed that over accumulation of free amino acida such as Arg and Ala may inhibit spike growth in the *gs1.1-1* mutant.

In summary, the three *TaGS1* genes differentially responded to N availability. *TaGS1.1* is up-regulated by low-N and is the major *GS1* gene expressed in roots and shoots at transcriptional and protein levels, supporting the importance of this gene in NH_4_^+^ assimilation and wheat adaptation to low-N stress. *TaGS1.1* has essential roles in remobilization, and its mutation delays leaf N loss and senescence during grain filling. Lack of *TaGS1.1* causes growth deficiency in an organ age-dependent manner and reduces yield by impairing spikelet and grain number. The current study and the reported literature showed that GS1.1 orthologues in wheat, rice, and maize were functionally diversified in adaptation to N resources and in mediating yield component formation. As such, strategies for the use of *GS1* genes in increasing yield and NUE should be optimized in different crops.

## Acknowledgments

This research was supported by The National Key Research and Development Program of China (2016YFD0100706) and the Strategic Priority Research Program of the Chinese Academy of Sciences (Precision Seed Design and Breeding, Grant No. XDA24010202).

## Author contributions

YZW and WT performed the experiments; YPW and CXG performed genome editing; WT and XO identified the *gs1.1* mutant; XH and XQZ assisted field experiments; WT and YPT wrote the article.

## Supplemental Data

**Supplementary Document S1.** Off-target prediction

**Supplemental Figure S1.** Phylogenetic analysis of GS proteins in plants.

**Supplemental Figure S2.** Expression analysis of *GS1* genes in different organs of wheat.

**Supplemental Figure S3.** Preliminary measurement of agronomic traits in the wild type KN199 and the *gs1.1* mutants.

**Supplemental Figure S4. The difference of** heading time between the wild type and *gs1.1-1* mutant plants.

**Supplemental Figure S5.** Dry weights of the aerial parts in the main culm at different developmental stages in the 2017-2018 growing season.

**Supplemental Figure S6.** The expression of N assimilation genes in the WT and *gs1.1-1* mutant seedlings.

**Supplemental Figure S7.** The expression of *GS* and *GOGAT* genes in flag leaves of the WT and *gs1.1-1* mutant plants under high-N conditions in the field experiment of the 2017-2018 growing season.

**Supplemental Figure S8.** Concentration of free amino acids in young spikes and mature seeds in the field experiment of the 2017-2018 growing season.

**Supplemental Figure S9.** N concentrations in different aerial parts of the main culm at different developmental stages in the field experiment in the 2017-2018 growing season.

**Supplemental Table S1.** PCR primers used in this study.

**Supplemental Table S2.** Gene Ids and former names of *GS* genes in wheat.

**Supplemental Table S3.** Agronomic traits of the wild type and *gs1.1-1* mutant under low-N and high-N conditions in the field experiment of the 2018-2019 growing season.

**Supplemental Table S4.** Concentration of N metabolites in shoots and roots of the wild type and *gs1.1-1* mutant grown under SN, LN, and AN conditions.

